# Microbes and diet reshape the intestine via distinct cellular dynamics

**DOI:** 10.1101/2025.09.17.676798

**Authors:** Alessandro Bonfini, Nicolas Buchon

## Abstract

**Background:** The intestine adapts to environmental stimuli by dynamically altering its size. Infections causes shrinkage and subsequent regrowth to original intestinal dimensions, suggesting homeostasis, whereas diet can induce adaptive changes. Whether diet and infection differ in the kinetics, magnitude, or cellular mechanisms that drive intestinal resizing, and whether intestines use a fixed size “memory” versus an adaption to nutrient availability, still remains unclear.

**Objective:** To determine whether intestinal regrowth after infection reflects a fixed “memory” of a target size (homeostasis) or an adaptation to the nutritional state during regrowth, and to identify the cellular modalities that drive infection- and diet-induced resizing.

**Design:** Using *Drosophila melanogaster* as a model, we quantified intestinal size, cell size, cell number, and epithelial turnover in response to dietary shifts and oral bacterial infections, both individually and in combination.

**Results:** Infection-induced atrophy occurred only when the intestine was initially large, and regrowth required nutrient availability. Distinct cellular mechanisms underpinned changes in intestinal size: diet-induced growth in unchallenged intestines was mediated by enterocyte hypertrophy, whereas post-infection regrowth required intestinal stem cell (ISC) proliferation. Consistently, ISC ablation impaired regrowth only after infection. Bacteria functioned both as pathogens and nutrients, triggering intestinal shrinkage when damaging a large organ, but promoting growth when consumed by nutrient-deprived flies.

**Conclusion:** Intestinal regrowth after infection reflects a context-dependent, diet-driven adaptation, not a fixed intestinal size memory. Diet and infection engage distinct, stimulus-specific cellular programs to reshape the midgut, revealing a modular logic of organ plasticity with implications for tissue repair and metabolic regulation.

**Summary box:** *What is already known on this topic:*

- Intestinal size is regulated by epithelial turnover and responds dynamically to dietary and microbial stimuli.

*What this study adds:*

- Post-infection intestinal regrowth is governed by dietary input rather than an intrinsic memory of pre-infection intestinal size.
- Intestinal resizing is driven by distinct cellular programs: diet-induced intestinal growth occurs independently of stem cells, while post-infection intestinal regrowth requires stem cell proliferation.
- Diet, rather than microbes, is the primary driver of intestinal size, while pathogens influence the mode of size change by increasing cellular turnover.
- Pathogenic microbes act as both sources of damage and nutrients, influencing organ resizing.

*How this study might affect research, practice, or policy:*

- Reframes intestinal regeneration as a flexible, context-dependent adaptation rather than a fixed homeostatic process.
- Identifies nutritional status as a critical determinant of recovery following intestinal injury.
- Offers a conceptual framework for understanding epithelial plasticity in health and disease, emphasizing how the balance of cell size and stem cell activity coordinates adult intestinal size.

## Introduction

Since antiquity, scholars have sought to explain how living organisms maintain internal stability in the face of external fluctuations. In the nineteenth century, Claude Bernard introduced the concept of the *milieu intérieur* [1], which Walter B. Cannon later reframed as “homeostasis”: the active maintenance of a relatively constant internal state [2]. This concept implicitly assumes a single, fixed set point for physiological equilibrium. In recent decades, the notion of homeostasis has been extended beyond physiological parameters to include structural and functional integrity at the level of tissues and organs, particularly those maintained by stem cell activity.

Organ size is regulated by dynamic processes that balance cell production, loss, and growth [3,4]. In adult epithelia such as the intestinal lining, the entire epithelium renews every 3-5 days, with approximately 300 million cells replaced daily in humans [5]. This renewal process is tightly regulated to support not only tissue maintenance but also adaptive remodeling in response to environmental cues, including diet and microbial exposure [6]. While both infections and nutritional changes are known to reshape the intestine [3,4,7,8], it remains unclear whether they do so via shared or distinct cellular programs, and how these signals might interact to define organ size.

The adult *Drosophila melanogaster* midgut is a well-established model for studying these mechanisms due to its genetic tractability and its conservation of key regulatory pathways and cellular architecture with the mammalian intestine [9]. The midgut epithelium primarily consists of polyploid enterocytes (ECs), which are continuously replenished by intestinal stem cells (ISCs) [10–12]. ISCs generate either enteroblasts (EBs), which differentiate into ECs, or pre-enteroendocrine (pre-EE) cells, which mature into EEs [13]. A powerful genetic toolkit enables precise manipulation of gene function, dietary inputs, and microbial exposures in a temporally and cell type– specific manner [14–16]. Additionally, clonal analysis systems allow quantification of tissue turnover rates in the gut, where all cells can be measured across the entire organ, enabling multiscale investigation of cellular and tissue-level dynamics.

In this context, homeostasis refers to an organ’s ability to maintain its structure, function, and size despite environmental challenges. This is achieved through tightly regulated epithelial turnover, which prevents abnormal atrophy or overgrowth in healthy animals. In the Drosophila midgut, turnover is maintained via steady-state feedback mechanisms that match EC loss with ISC-driven replacement [4]. This process is resilient, allowing the intestine to respond to acute damage through compensatory stem cell-driven regeneration [17–20]. For instance, oral infection with pathogenic bacteria such as *Erwinia carotovora* (*Ecc15*) [21] leads to delamination and death of up to half of midgut ECs, yet ISC proliferation and EB differentiation restore tissue architecture and size within 48 hours [7]. This regenerative response is evolutionarily conserved: various infections cause epithelial remodeling across species. In mice, *Rotavirus*, *Cryptosporidium*, and *Trichinella spiralis* induce villus blunting and crypt hyperplasia [22–24], while *Giardia lamblia* in gerbils similarly shortens villi and deepens crypts [25]. Comparable effects are seen in rats [26], sheep [27] and piglets [28]. In humans, HIV enteropathy leads to villus atrophy and crypt hyperplasia, contributing to diarrhea, wasting, and malabsorption [29]. Moreover, individuals with repeated enteric infections show shortened villi and enlarged crypts on biopsy [30].

Diet also induces substantial, reversible changes in midgut size. Nutrient-rich diets promote midgut expansion via increased ISC proliferation and EC hypertrophy, whereas nutrient-poor or imbalanced diets reduce midgut size by decreasing cell size and increasing cell loss. Specifically, protein-rich diets drive net tissue expansion through increased symmetric ISC divisions [8] and EC enlargement [3], while low-protein, high-sugar diets promote intestinal atrophy through EC autophagy and cell loss [3]. Like infection-induced remodeling, diet-driven intestinal plasticity is evolutionarily conserved. In mice, starvation causes intestinal atrophy [31], high-fat diets reduce villus length [32] and Western diets increase the villus-to-crypt ratio, expanding the absorptive surface [33]. In humans, malnutrition results in villus blunting and crypt hyperplasia [30], a phenotype that can be reversed upon nutritional rehabilitation [34].

At first glance, these two models appear contradictory. Post-Infection regeneration is often framed as a return to a fixed, pre-injury midgut size, whereas diet elicits a flexible adaptation in midgut size. Reconciling these paradigms raises a key question: does the midgut regrow to a homeostatic set point after damage, or is midgut size recalibrated based on post-infection dietary conditions?

Adding further complexity is the role of host–microbe interactions. While bacteria such as *Ecc15* [21], *Pseudomonas entomophila* [35] and *Serratia marcescens* [36] cause midgut damage, commensal microbes offer benefits, including colonization resistance against pathogens [37]. Moreover, the microbiota supports Drosophila development [38,39] and influence nutrient metabolism, absorption, and potentially midgut epithelial renewal [40]. Notably, under dietary stress, midgut microbes can serve as nutritional resources [41]. This dual role raises the question: are pathogenic microbes purely damaging agents, or do they also contribute to host nutrition, and how might this duality shape intestinal responses?

To address these questions, we combined defined dietary regimens with controlled bacterial infections in Drosophila and recorded the cellular basis of midgut size changes at a multiscale level. By measuring stem cell activity (cell gain), total cell number (net balance of gain and loss), and cell size, we found that post-infection intestinal regrowth does not return the midgut to a fixed set point, but instead reflects a flexible, diet-dependent adaptation process. Distinct cellular mechanisms underlie this plasticity: stem cell activity drives infection-induced changes in midgut size but is dispensable for diet-driven resizing, which primarily relies on EC hypertrophy and atrophy. These findings reveal a modular logic of midgut remodeling and provide a framework for understanding how epithelial tissues integrate concurrent environmental signals. Given the importance of stem cell renewal and epithelial dynamics in human intestinal disorders, this work offers insights into how the midgut balances repair, function, and adaptation, and may inform new therapeutic strategies for modulating tissue plasticity in health and disease.

## Materials and methods

See supplementary materials and methods.

## Results

### Infection reduces midgut size in nutrient-rich but not nutrient-poor conditions

To compare how diet and infection influence midgut epithelial turnover and size, we designed a matrix of experimental conditions integrating dietary shifts and microbial exposure (online supplemental figure 1A). All flies were reared on a common standard diet. At eclosion, they were transferred to one of two diets: a high-yeast (HY), protein-rich diet, which supports large midgut growth, or a high-sugar (HS) diet, which maintains a smaller midgut, consistent with previous findings [3,42]. Each dietary group was then either infected with the bacterial pathogen *Erwinia carotovora carotovora* 15 (*Ecc15*) or left uninfected. After eight hours, each group was moved to one of three dietary conditions: either HS, HY, or starvation (a non-nutritive agar diet to mimic nutrient deprivation). This design produced a matrix of twelve conditions representing all combinations of diet and infection history. To monitor the full progression across conditions, flies were dissected before infection and at 12, 24, 48, 96, and 240 hours after the initial infection.

On the HY diet, *Ecc15* infection led, as expected, to a marked and rapid reduction in midgut size within 12 hours, followed by progressive regrowth until 48 hours post infection (figure 1A–C; quantification in figure 1D, E). In contrast, flies maintained on the HS diet, whose midguts were already smaller at the time of infection, did not exhibit further shrinkage (figure 1F–H; quantification in figure 1I, J). Instead, we observed a modest increase in midgut size, peaking at 24 hours following initial *Ecc15* exposure. These results indicate that infection-induced midgut shrinkage is not a universal phenomenon and is contingent upon an initially large midgut deriving from a nutrient-rich context, underscoring the importance of dietary state in shaping midgut responses to microbial challenges.

**Figure 1:**
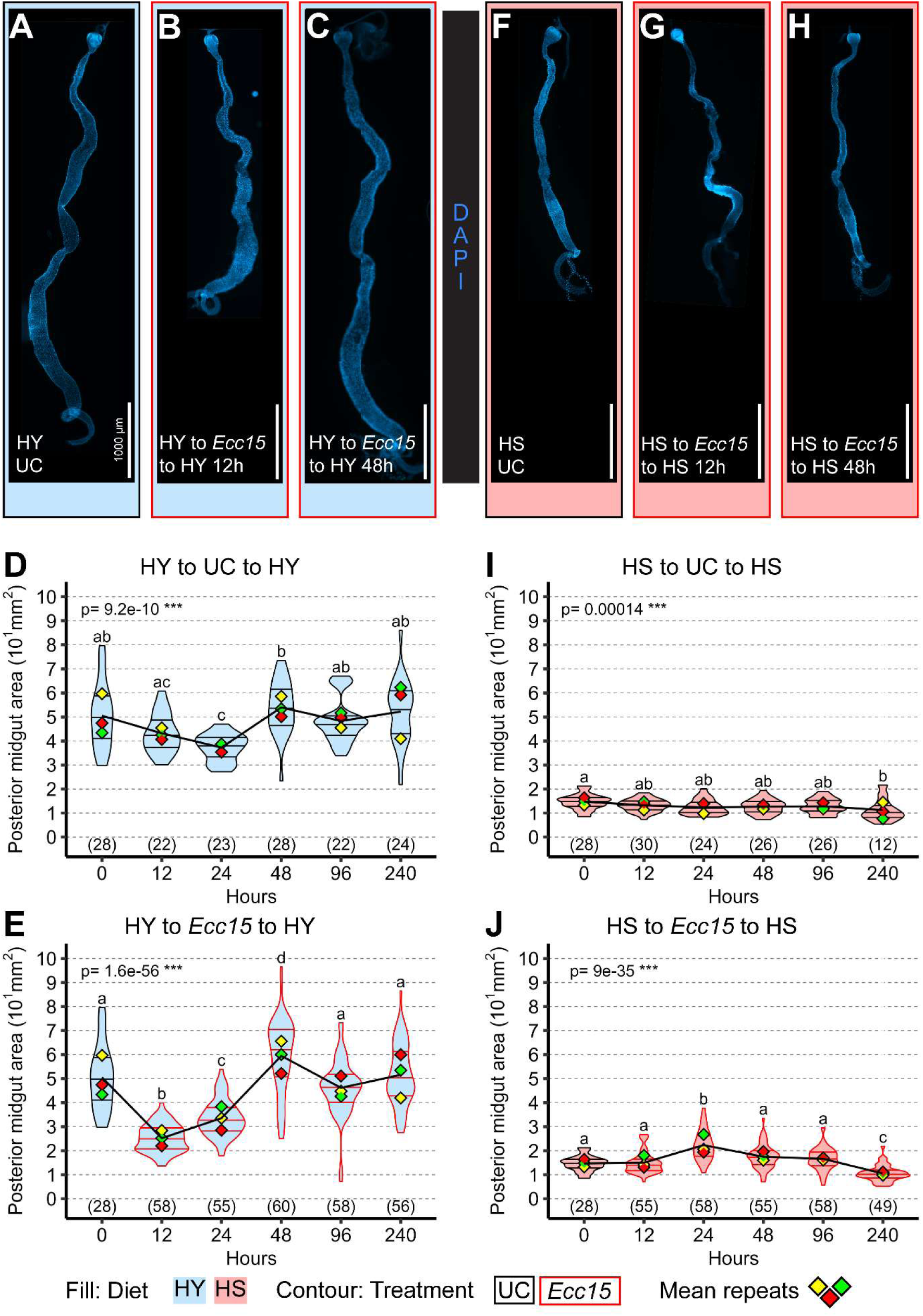
Infection does not systematically trigger midgut shrinkage. (A-E) Midguts constantly on high yeast (HY) diet have a stable size, as shown in (A) and quantified (D) for the posterior midgut. Upon infection, midguts initially shrink (B) and then regrow (C), as quantified (E) for the posterior midgut. (F-J) Midguts constantly on high sugar (HS) diet also have a stable size (F), as quantified (I), but smaller than midguts on HY (A, D). When infected, midguts that are already small due to HS diet do not further shrink (G), as quantified (J), but transiently enlarge at 24h post infection (J) to then return to their initial size (H). Images are 10x composite tiling of midguts stained with DAPI. Scale bars are 1000 µm for all images. For the violin/dot plots shown in this figure, colored lozenges represent means of replicate experiments. Black line connects total mean of each sample to show timeline of changes. Violin plot fills are color-coded according to diets (HS = red, HY = light blue) and their contour indicates treatment (UC = black, *Ecc15* = red). Numbers in parentheses at the bottom of charts indicate sample sizes. p indicates the result of ANOVA for samples in a single chart, and groups are obtained with Tukey Post Hoc test (letters above violins).

### Post-infection intestinal regrowth depends on dietary context

We next asked whether midgut regrowth after infection reflects restoration to a pre-infection set point, or whether it is shaped by the current nutritional environment. Flies shifted to HS without infection exhibited a progressive reduction in posterior midgut size under unchallenged conditions (figure 2A, B, D), as previously reported [3]. When infected prior to the shift to HS, midguts first underwent infection-induced shrinkage (figure 2C, E). While shifting flies back to HY (figure 1E) supported midgut regrowth after infection, shifting flies to HS failed to promote midgut regrowth (figure 2E). Similar results were obtained when shifting flies to starvation conditions (Ag, online supplemental figure 2A-B). Conversely, HS-to-HY flies showed robust midgut growth regardless of infection, suggesting that dietary nutrients, not infection history, determine final midgut size. These findings indicate that midgut regrowth is not a return to a fixed homeostatic size, but rather an adaptation to prevailing nutritional conditions.

**Figure 2:**
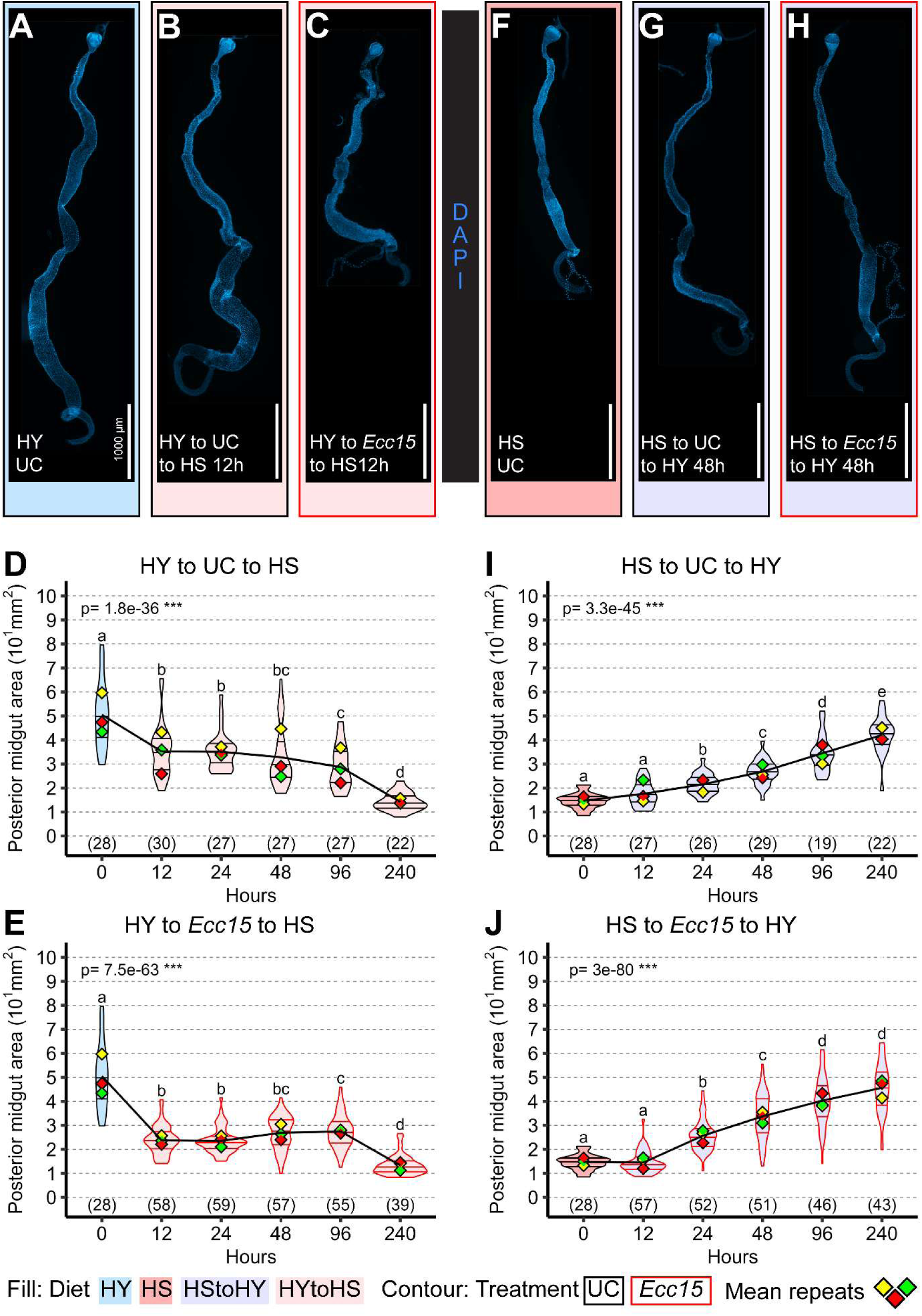
Nutrients are required for post infection organ regrowth. (A-E) Midguts shifted from high yeast (HY, A) diet to high sugar diet (HS, B) in unchallenged conditions (UC) progressively decrease in size (D). Upon infection, midguts shrink, and then maintain this small size (C), as quantified (E) for the posterior midgut. (F-J). Midguts shifted from HS (F) to HY (G) in UC conditions progressively increase in size (I). When infected, midguts also progressively increase in size (H), with a similar trajectory (J) as UC midguts (I). Images are 10x composite tiling of midguts stained with DAPI. Scale bars are 1000 µm for all images. For the violin/dot plots shown in this figure, colored lozenges represent means of replicate experiments. Black line connects total means of each sample to show timeline of changes. Violin plot fills are color-coded according to diets (HS = red, HY = light blue, HY to HS = pink, HS to HY = purple) and their contour indicates treatment (UC = black, *Ecc15* = red). Numbers in parentheses at the bottom of charts indicate sample sizes. p indicates the result of ANOVA for samples in a single chart, and groups are obtained with Tukey Post Hoc test (letters above violins). Panels A and F, and time 0 in the charts, are the same as in figure 1, and are presented here again for clarity. This data was derived from a single large experiment, as shown in online supplemental figure 1.

### Midgut resizing following diet or infection emerges through different cell dynamics

Our data show that both infection and diet can lead to midgut shrinkage, and that diet dictates the extent of midgut resizing. However, whether similar midgut sizes arise from comparable cellular processes remains unclear. To address this, we used a lineage-tracing system previously developed by our group [3] that simultaneously quantifies EC loss and gain. Through a pulse-chase–like design, ECs transiently express a stable His2B-RFP fluorophore, labeling ECs present at the initial time point but not newly generated ones. Over time, labeled (RFP⁺) ECs are progressively lost and replaced by newly generated, unlabeled (RFP⁻) ECs. Tracking the decline of RFP⁺ ECs and the rise of RFP⁻ ECs allows us to infer rates of epithelial cell loss and renewal.

We applied this assay across all twelve diet/infection conditions, tracking EC labeling over six time points spanning ten days (representative images for 48 hours in figure 3A-H, complete dataset for all conditions is shown in online supplemental figure 3 and online supplemental figure 4). From these data, we first calculated the rate of cell loss or gain per hour at each time point, inferred from the change in cell numbers relative to the previous time point divided by the elapsed time (online supplemental figure 4A). This allowed us to visualize the timing of maximal cell loss and gain associated with infection, as well as the overall trajectory of cell dynamics. We then summarized these patterns by calculating absolute rates of cell loss at 12 hours (figure 3I), cell loss at 240 hours (figure 3J), and cell gain (figure 3K). Because flies raised on HS and HY diets start with different midgut sizes, we also computed proportional rates, cell loss or gain per initial total cell number, and derived a replacement ratio to estimate the coupling between loss and renewal (figure 3L).

**Figure 3:**
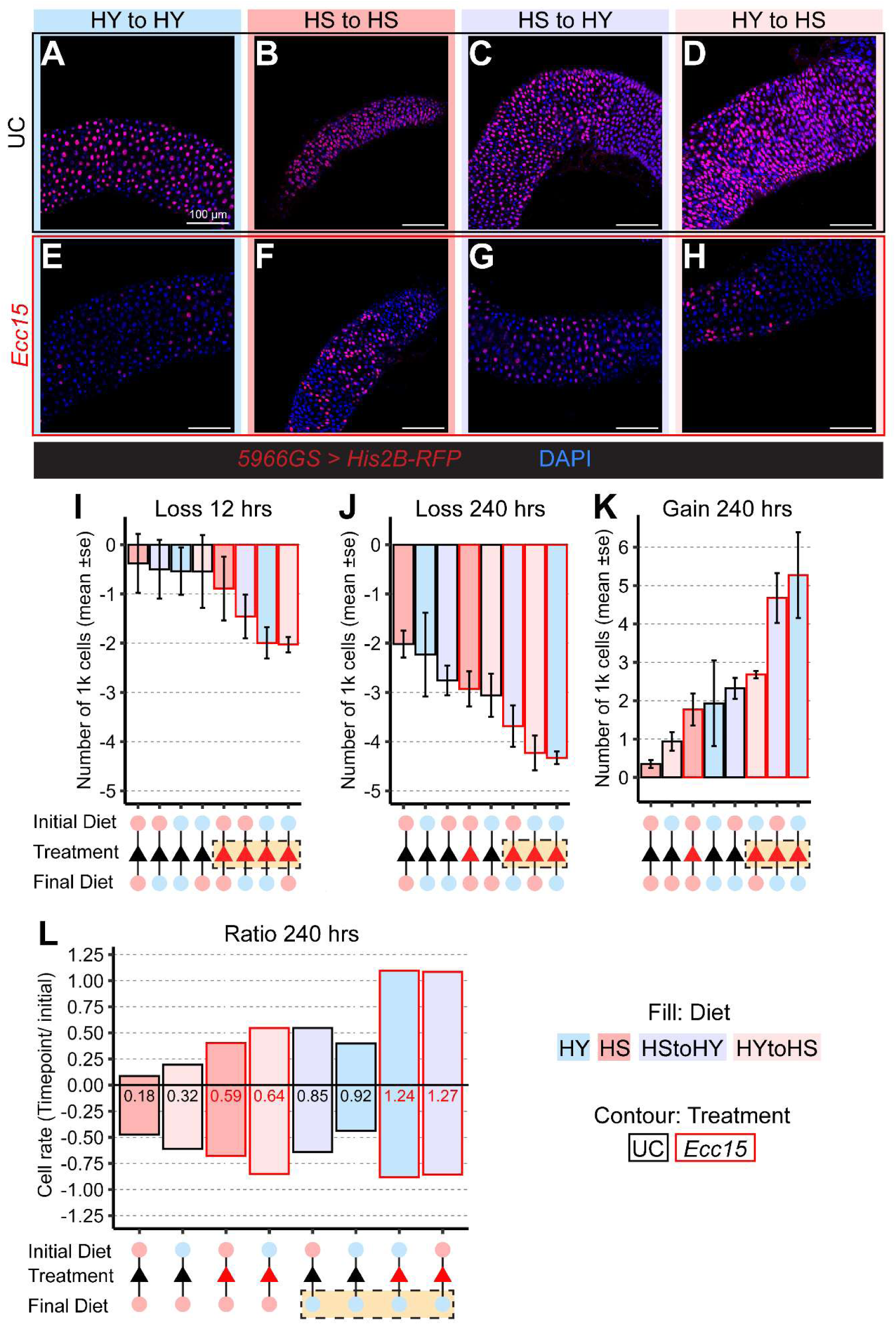
Infection dominates loss and gain while diet decides overall ratio. A-H) Representative pictures at 48 hours post infection of the region 4 of *5966GS>His2B-RFP* midguts unchallenged (A-D) or challenged with *Ecc15* (E-F). I-L) Infection with *Ecc15* overall results in increased amplitude of both cell loss (I, J) and cell gain (K), but overall growth (Ratio of gain/loss) is found higher on samples shifted after treatment on HY diet (L). Images are 20x maximum projection of midguts expressing His2B-RFP (red) and stained with DAPI (blue). Scale bars are 100 µm for all images. Bar plot fills are color-coded according to diets (HS = red, HY = light blue, HY to HS = pink, HS to HY = purple) and their contour indicates treatment (UC = black, *Ecc15* = red).

Overall, our data reveal that infection and diet influence epithelial dynamics through distinct mechanisms. Infection with *Ecc15* triggered an acute spike in EC loss [7], reflected in a consistent increase in the absolute rate of cell loss (figure 3I). Infected samples continued to show the highest levels of cell loss even at later time points. At these later stages, however, dietary effects also became apparent, with midguts maintained on the HS diet exhibiting comparable levels of loss, particularly when small midguts were both infected and kept on HS diet (figure 3J). Conversely, cell gain (figure 3K) was generally elevated under infection conditions and further enhanced by a shift to the HY diet. In contrast, shifts to the HS diet were associated with reduced cell gain and slower, progressive cell loss. Importantly, the rates of cell loss and gain did not consistently coincide across treatments, underscoring a partial decoupling between epithelial loss and renewal dynamics.

Replacement ratio analysis further underscored these differences. High replacement ratios (more cells made than lost) occurred exclusively in flies shifted to the HY diet, regardless of infection status. In contrast, on the HS diet, whether infected or not, replacement ratios remained below one (more cells lost than made), indicating incomplete replacement of lost ECs. The only condition approaching a replacement ratio of one was uninfected flies maintained continuously on the HY diet, which corresponds to a stable, large midgut size and reflects apparent homeostatic turnover. Infection on the HY diet elevated cell gain above cell loss, consistent with damage-induced regeneration (figure 3L).

These observations were further supported by raw cell count trends. Midguts from HS-fed flies showed a gradual reduction in total EC number and minimal turnover (online supplemental figure 3C). In contrast, flies reared on HS, then infected and returned to HS, exhibited increased epithelial turnover (online supplemental figure 3D), despite having similar final cell numbers and midgut size (figure 1I, J). Similar trends were observed in flies fed on HS and shifted to starvation diets (online supplemental figure 3K, L). Altogether, our findings demonstrate that bacteria and diet independently regulate distinct aspects of epithelial dynamics, and that rates of EC loss and gain are differentially modulated by infection-induced damage and nutritional state.

### Diet and bacteria impact ISC proliferation and enterocyte size differently to change midgut size

To further understand how diet and damage alter the mechanisms of resizing, we focused on two conditions with an identical starting point and that result in similarly sized midguts but differ in cellular dynamics. When flies transition from the HS to the HY diet, the midgut expands over several days, with or without infection (figure 4A). However, as shown in online supplemental figure 3 and further illustrated in figure 4B-C and quantified in figure 4D-E, this identical midgut size in the presence of infection is associated with higher epithelial turnover, resulting in a significantly greater final number of ECs in infected samples compared to uninfected ones, indicating that midgut size is achieved differently depending on infection status.

**Figure 4:**
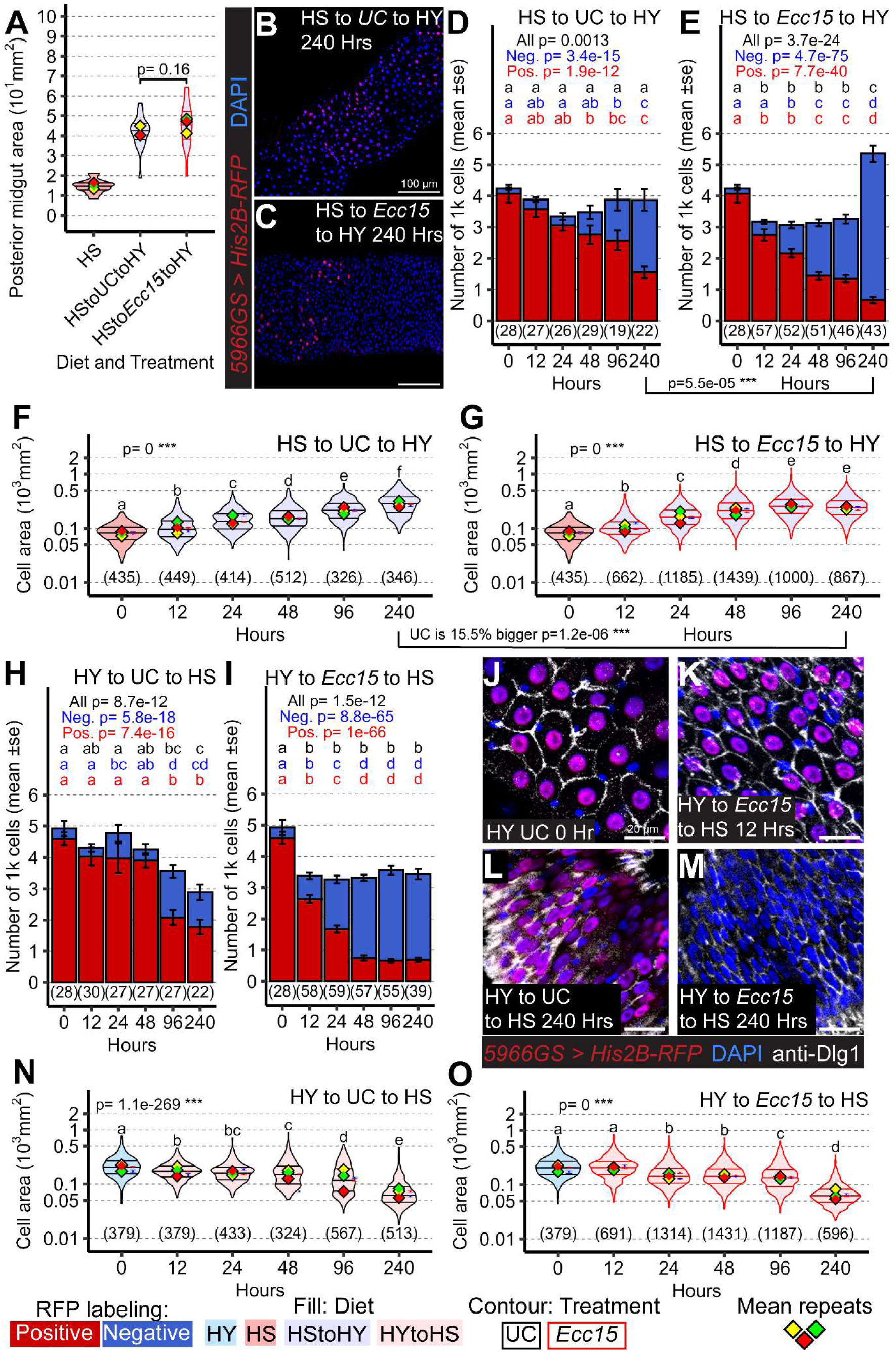
Midgut size change following diet or infection emerges through different cell dynamics. A-G) Midguts of flies initially on HS diet and then shifted to HY with or without infection grow to similar size after 240 hours (A). However, the number of cells on HS to UC to HY samples at 240 hours (B, D) is sig. lower than the one from HS to *Ecc15* to HY samples (C, E). This difference is reflected also in the size of differentiated cells (F, G). H-O) Midguts originally on HY and then shifted to HS show that loss of midgut size upon infection is through loss of cells (quantified in H, I), and not reduce size of cells (examples in J-M, and quantified in N, O) while nutrition leads to both loss of cells (H 240 hours) and cell size (N 240 hours). Images are 20x maximum projection of midguts expressing His2B-RFP (red) and stained with DAPI (blue) and anti-Dlg1 (white). Scale bars are 100 µm for B and C, and 20 µm for J-M. For the violin/dot plots shown in this figure, colored lozenges represent means of replicate experiments. Black line connects total means of each sample to show timeline of changes. Violin plot fills are color-coded according to diets (HS = red, HY = light blue, HY to HS = pink, HS to HY = purple) and their contour indicates treatment (UC = black, *Ecc15* = red). Numbers in parentheses at the bottom of charts indicate sample sizes. p indicates the result of ANOVA for samples in a single chart, and groups are obtained with Tukey Post Hoc test (letters above violins). 1 on 1 comparison across diets were performed independently for the specific time points reported. Charts showing for each time point the amount of RFP positive (red) and RFP negative cells (blue). Statistics refers to comparison in between same treatment, and compared for both total number of cells, RFP positive cells (old cells) and RFP negative cells (new cells) (D, E). p value external to samples connecting D and E, and F and G are to compare samples from different treatments. Panels D, E, H, I are the same as in online supplemental figure 3G, H, E, F. They are presented here again for clarity and included in the online supplemental figure 3 to offer a complete comparison.

Midgut size in Drosophila reflects both the number and size of ECs, which are independently regulated by diet [3]. We hypothesized that the relative contributions of proliferation and EC growth may differ between infection- and diet-driven responses. To test this, we measured EC area in the same samples used for turnover analysis. As expected, EC size increased over time in both infected and uninfected flies shifted from HS to HY (figure 4F-G). However, by 10 days post-infection, ECs were significantly larger in unchallenged controls (15.5% percent larger than *Ecc15* treated midguts at the same timepoint), conditions in which turnover was lower, indicating that post infection midgut regrowth favors proliferation over hypertrophic expansion of EC size.

Midgut shrinkage could also be influenced via different mechanisms in response to bacteria and diet. Midgut shrinkage occurs when flies reared on a HY diet are exposed to *Ecc15* or switched to the HS or starvation diet (figure 2D-E, online supplemental figure 2B). Despite producing similar reductions in midgut size, albeit with different kinetics, these two conditions resulted in different degrees of cell loss: greater in infected flies (figure 4I) than in flies experiencing a dietary shift alone (figure 4H). To further investigate how midgut shrinkage occurs in each case, we monitored EC size. A shift from a large midgut initially on HY diet to the HS diet caused a progressive reduction in EC size over time (figure 4J, L, N). In contrast, infection, despite shrinking large midguts already at 12 hours, did not reduce EC size (figure 4K, O, online supplemental figure 5A, B), with cell size declining later on as per unchallenged samples (figure 4M, O). Infected midguts originating from small HS diets even exhibited an increase in EC size (online supplemental figure 5D, F). These results indicate that infection-induced midgut shrinkage stems from cell loss rather than changes in cell size. Altogether, these findings demonstrate that bacterial infection and dietary restriction drive midgut size reduction through distinct cellular mechanisms: infection increases cell loss, while dietary cues primarily reduce EC size.

### Stem cells are required for regrowth after infection but dispensable for diet-induced growth

Our results suggest that ISC proliferation and EC hypertrophy contribute differently to midgut resizing, depending on whether the stimulus is dependent on dietary shifts or post-infection. Supporting this distinction, we previously showed that flies shifted from an HS to an HY diet can expand their midguts even when progenitor cells are genetically ablated, through compensatory EC hypertrophy [3].

In line with our current findings that proliferation is more prominent following infection than in response to dietary cues, we hypothesized that while ISCs and EB activity is not needed for naïve midguts, it may be essential for post-infection midgut resizing. To test this, we genetically ablated ISCs and EBs by overexpressing the pro-apoptotic gene *reaper* (*esg^TS^>rpr^OE^*). In these flies, the midgut expanded normally in response to a dietary shift (HS to UC to HY; figure 5B) but failed to regrow following infection. This failure was consistent across all dietary conditions, although in some cases it manifested as an inability to maintain midgut size rather than to initiate regrowth (figure 5A, C, D).

**Figure 5:**
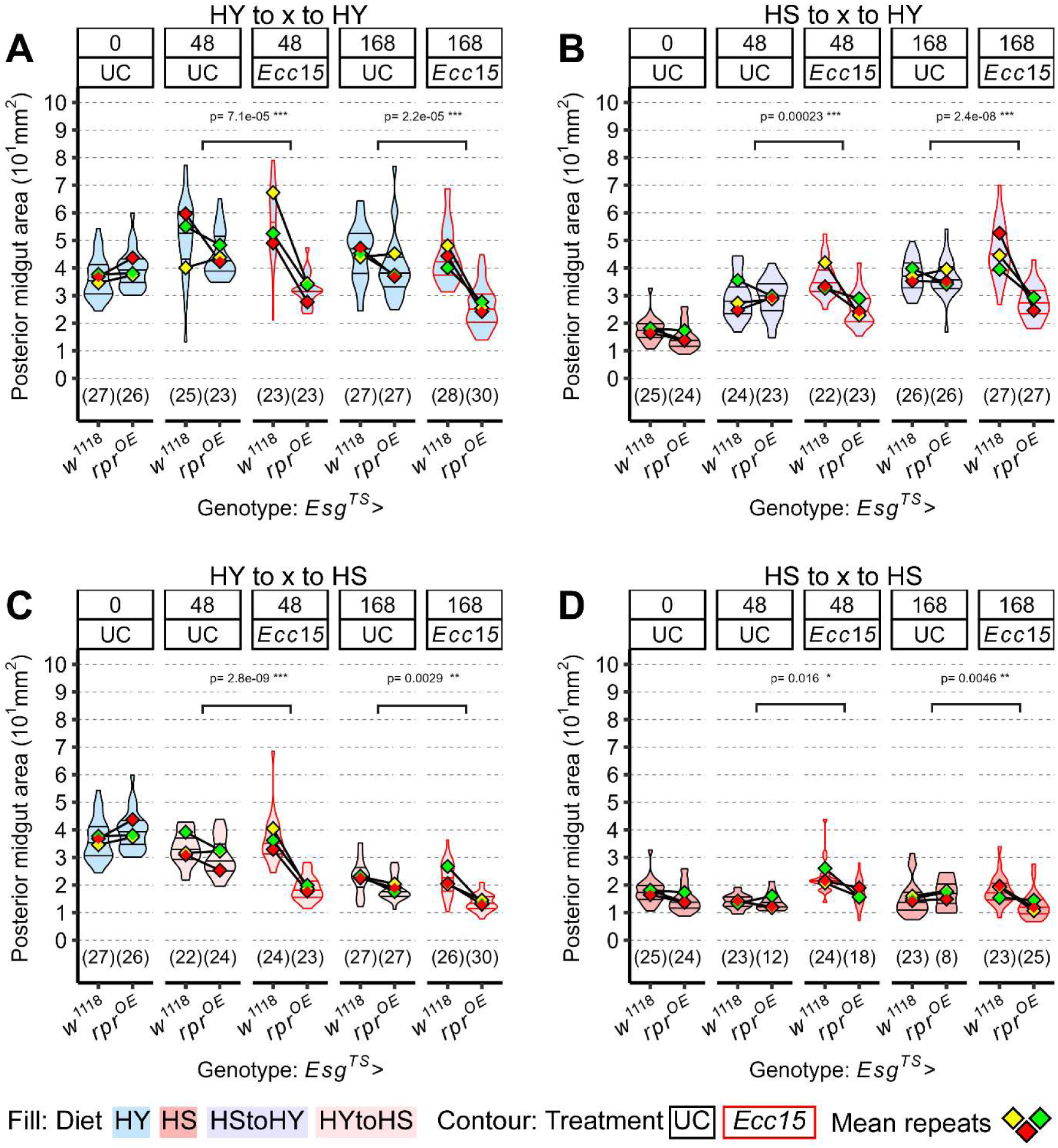
Stem cells are required for growth only post infection. A-D) *Esg^TS^>Rpr^OE^* midguts devoid of progenitor cells are still able to keep their size/ grow upon being on HY diet, but cannot do so following infection (A, B), resulting in an overall smaller size of the midgut compared to control midgut *Esg^TS^ x w^1118^*. This effect is visible also for midguts shifted to HS diet following infection, with *Esg^TS^>Rpr^OE^* being generally smaller than their control counterpart when infected (C, D). For the violin/dot plots shown in this figure, colored lozenges represent means of replicate experiments. Black line connects total means of each sample to show timeline of changes. Violin plot fills are color-coded according to diets (HS = red, HY = light blue, HY to HS = pink, HS to HY = purple) and their contour indicates treatment (UC = black, *Ecc15* = red). Statistical tests are for the interaction between genotype and treatment.

These findings indicate that progenitor activity is dispensable for diet-induced growth, which primarily relies on enterocyte hypertrophy. In contrast, ISC activity is essential for infection-triggered regeneration, underscoring a context-specific requirement for stem cells in intestinal resizing.

### The impact of bacteria on midgut size and host survival is diet-dependent

As mentioned in the introduction, microbes can be both a source of damage and an essential nutrient source for the host. We first established that, in our system, non-pathogenic bacteria can indeed act as nutrients and visibly influence midgut size (figure 6). We compared the evolution of midgut size over seven days in flies maintained on the protein-poor HS diet, either alone or with repeated feedings (figure 6A) of the non-pathogenic microbe *E. coli*. For repeated feeding, flies were exposed to bacteria for 8 hours and then shifted back to HS diet for 16 hours before the next bacterial exposure. Interestingly, ingestion of *E. coli* led to a slight but significant increase in midgut size over this seven-day period (figure 6B–C), consistent with previous findings suggesting that bacteria can act as nutrients that promote midgut growth.

**Figure 6:**
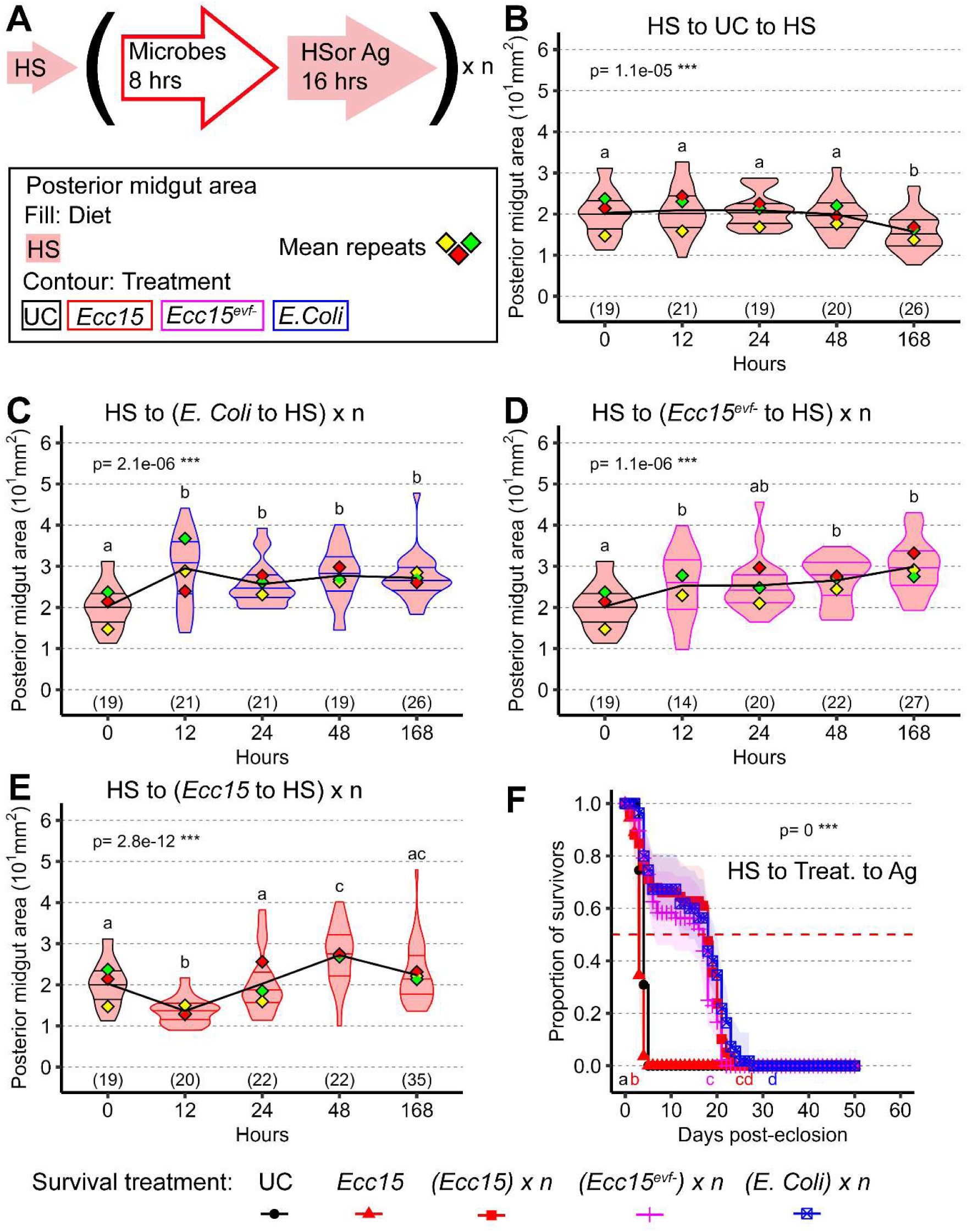
Diet reveals the nutritive and pathogenic dual effect of bacteria. A) Scheme depicting timeline of experiments with repeated treatments in this figure and online supplemental figure 6. For the violin/dot plots shown in this figure, white dots represent single posterior midgut area measurements. B-E) Midguts of flies constantly on HS diet keep a relatively small midgut size, with further shrinkage upon prolonged stay on HS diet (B). Repeated addition of nonpathogenic microbes (*E. coli*, C and *Ecc15^evf-^*, D) have an immediate effect in increasing the size of the midgut. The same effect is visible with a slight delay even when adding the pathogenic *Ecc15* (E), which shows an increase in midgut size after an initial decline. F) Pathogenic bacteria like *Ecc15* can prove to be beneficial when supplemented to completely starving flies (HS to treat. to Ag), prolonging the lifespan of the flies compared to starvation or a single treatment of *Ecc15*. For the violin/dot plots shown in this figure, colored lozenges represent means of replicate experiments. Black line connects total means of each sample to show timeline of changes. Violin plot fills are color-coded according to diets (HS = red, HY = light blue, HY to HS = pink, HS to HY = purple) and their contour indicates treatment (UC = black, *Ecc15* = red). Numbers in parentheses at the bottom of charts indicate sample sizes. p indicates the result of ANOVA for samples in a single chart, and groups are obtained with Tukey Post Hoc test (letters above violins).

We next asked whether a bacterial pathogen such as *Ecc15* could also promote midgut growth. Demonstrating this is challenging, because *Ecc15* induces high levels of stress and epithelial damage, which normally lead to midgut shrinkage [7]. To address this, we used two strategies. First, we repeatedly exposed HS-fed flies to the avirulent strain *Ecc15Δevf*, which does not trigger epithelial damage [43]. Repeated ingestion of Ecc15Δevf promoted midgut growth on the otherwise non-stimulatory HS diet, similar to *E. coli* (figure 6D). Second, we examined the effect of repeated *Ecc15* infection in flies previously maintained on HS, a condition shown to prevent infection-induced midgut shrinkage (figure 2J). Under these conditions, repeated *Ecc15* ingestion did not lead to midgut shrinkage but instead resulted in a small, sustained increase in midgut size starting at 48 hours post-infection compared to uninfected HS controls (figure 6E). These treatments did not affect fly survival, with the exception of multiple *Ecc15* infections, which led to a small decrease (online supplemental figure 6A).

We previously showed that sugar in the HS diet can act as a mild stressor that antagonizes yeast-induced midgut growth [3]. Thus, an alternative explanation is that bacteria promote midgut growth by reducing sugar load rather than providing direct nutritional benefits. To test this, we repeated the *E. coli* and *Ecc15* feeding experiments but transitioned flies to a completely non-nutritive agar-based diet after bacterial exposure. This diet does not support midgut growth and leads to death within a few days (between two- and four-days post-transition; online supplemental figure 6B, C). Notably, repeated feeding with *E. coli*, *Ecc15Δevf*, or *Ecc15* still led to a small increase in midgut size (online supplemental figure 6D–F).

We next asked whether the nutritional value of microbes could provide measurable physiological benefits beyond their effects on gut size. Flies shifted from HS to the non-nutritive diet died rapidly in the absence of microbial supplementation, but strikingly, flies that received repeated bacterial feedings, including with pathogenic *Ecc15,* survived significantly longer (figure 6F). These findings demonstrate a dual role for pathogenic bacteria in the midgut: while they can induce epithelial stress and cell loss in nutrient-rich conditions, they also act as nutrient sources that promote organ growth and enhance survival under dietary restrictions.

## Discussion

Our findings demonstrate that intestinal size regulation in Drosophila operates through distinct cellular programs depending on whether the stimulus is dietary or infectious. While both environmental inputs ultimately reshape the midgut, they do so via distinct mechanisms: diet primarily modulates enterocyte (EC) size, whereas infection drives ISC-mediated turnover and regeneration.

### The midgut is primarily an adaptive organ

From a conceptual standpoint, previous work has distinguished between two modes of intestinal size control: homeostasis, in which feedback mechanisms maintain the original state under physiological conditions or restore it following damage to preserve a defined midgut size [4,7]; and allostasis, in which midgut size set points are adjusted according to current nutritional status [3,8]. We asked whether post-infection organ regrowth represents a homeostatic reset to a prior size, or an adaptive regrowth toward a size dictated by diet. In the homeostatic scenario, the midgut would regrow post-infection to its original, pre-damage size. In contrast, an adaptive response would result in the midgut adopting a new size set point appropriate to prevailing dietary input, even if that size is smaller than the pre-infection state. Conceptually, homeostasis requires a defined set point (pre-stimulus midgut size), a sensor to detect deviation (e.g., dying cell signals), and mechanisms to restore the original state (e.g., ISC proliferation, EC growth). Our data supports the adaptive model: when infected flies were shifted to a nutrient-poor diet, the midgut failed to regrow and instead shrank to match the size typically associated with the HS diet (figure 2). Thus, although infection triggers a regenerative response, the final organ size is determined by the nutritional conditions post infection, not by a fixed memory of the original size.

### Beyond size: differential roles of nutrition and infection

Are similarly sized organs structurally equivalent? When flies are shifted from HS to HY, the midgut grows to similar sizes regardless of infection. Yet, by ten days post-infection, the number of cells is significantly higher in infected samples (figure 4). This suggests that infection alters how the midgut grows (e.g., more cells), while diet sets the final size. Other data support this distinction: for instance, infection increases the proportion of enteroendocrine (EE) cells during midgut regrowth [44]. Consistent with this division of labor, we previously showed that stem cells are dispensable for diet-induced resizing, which relies on EC hypertrophy [3]. Here, we find that stem cell ablation blocks midgut regrowth after infection, highlighting a functional separation between adaptation and regeneration. This has important implications for regenerative biology: therapies may need to differentially target proliferation or cell growth depending on whether tissue stress arises from damage or diet.

Despite these differences in cellular mechanisms, the midgut’s capacity to dynamically resize remains striking. That midgut size is so finely tuned to diet, despite major differences in underlying cell behaviors, underscores its evolutionary importance. Maintaining a self-renewing organ like the intestine is energetically costly. In environments characterized by cycles of feast and famine, the ability to resize quickly and efficiently may be critical.

### What is the role of enterocyte shedding?

Cell loss is an important component determining midgut size. We found that EC shedding in response to infection is not a universal phenomenon. Infection of small midguts (e.g., flies maintained on an HS diet) did not trigger further midgut shrinkage or significant cell loss in the first 12 hours (figure 1), with only mild loss and limited turnover observed thereafter (online supplemental figure 4A). This raises the question: is EC shedding an active, regulated tissue response, or simply a byproduct of pathogen damage?

One possibility is that epithelial fragility varies with EC size or metabolic state. ECs in HS midguts are smaller and may already be under mild stress [3], potentially making them more resilient to additional insults. Alternatively, biomechanical or cytoskeletal differences in smaller midguts might impair shedding mechanisms (e.g., cramps, junctional remodeling). In this light, EC loss is not strictly pathogen-driven but also influenced by nutritional history and tissue architecture. This suggests EC shedding may be a facultative response, selectively engaged depending on regenerative capacity or the cost-benefit balance of losing absorptive surface. Whether manipulating apoptotic or extrusion pathways in large midguts can reduce cell loss, and how this would affect survival or immunity, remains an intriguing question for future studies.

### Proliferative potential vs nutritional fulfilment of target size

Damage induces proliferation even in flies shifted to nutrient-poor diets, as shown by increased cell loss and replacement (figure 3). However, this regenerative activity does not translate into increased midgut size. While damage (via cytokine signaling) has the potential to promote midgut growth, this potential is realized only if dietary conditions support it. We suggest this is not simply due to limited resources, but reflects an active, context-sensitive decision: for example, flies infected with *Ecc15* and returned to an HS diet showed only a transient increase in midgut size, likely fueled by bacteria-derived nutrients, before shrinking again. This highlights the midgut’s tight coupling of intestinal growth dynamics to nutritional status.

We previously proposed the existence of a “size meter” in the context of midgut biology that calibrates intestinal size in response to diet, pushing the organ to reach a certain size regardless of the modality of growth or shrinkage [3]. We now observe that this concept also holds true across changes due to infective damage. To incorporate our newer finding, we propose that intestinal size is governed by “bounded plasticity”, that includes constraints by anatomical and physical limits: the maximum intestinal size may be bound by the space available in the body cavity, while the minimum is likely set by structural necessities, such as the mouth-to-anus distance. Within these boundaries, intestinal size is not fixed but fluctuates dynamically. Rather than diet determining a set-size to reach, it determines the direction of change (growth vs. shrinkage), while stress and damage modulate stem cell proliferation, enabling the tissue to realize or resist those changes. In this model homeostasis is an emergent property at the upper size boundary, where upon a growth inducing diet, cell loss is rapidly compensated to preserve maximum functional size. Furthermore, changes in intestinal size can arise from modulation of either cell number, influenced by damage and stem cell activity, or cell size, a more flexible and energetically efficient mode of adaptation. Cytokines, by modulating ISC proliferation post infection, change the contribution of cell number, and cell size plastically complete this process. Additionally, the proximity to the maximum and minimum intestinal size may be by itself a regulator of the speed of change in intestinal size. This framework unifies dietary and regenerative responses under a shared logic of constrained but highly responsive tissue remodeling.

### The dual nature of microbes

While pathogens like *Ecc15* induce epithelial stress and cell loss, they can also promote midgut growth and prolong survival under nutrient deprivation. Thus, microbes act as both stressors and resources, depending on physiological context. This duality may reflect an ecological strategy: ingestion of bacteria during nutrient scarcity provides nutritional benefits and enhances survival. Repeated *Ecc15* exposures led to progressively reduced damage and modest net growth, at a slight cost to survival (figure 6), suggesting a form of innate priming. Such priming may recalibrate ISC responsiveness or EC resilience over successive infections, reminiscent of trained immunity. This could enhance the midgut’s ability to extract nutrients from future microbial exposures, even pathogenic ones.

## Conclusion

In the broader context of midgut physiology and disease, these new findings offer insight into disorders marked by disrupted epithelial turnover, such as inflammatory bowel disease or malnutrition. Dissecting the distinct pathways that control stem cell– driven regeneration versus cell size–based growth could inform therapeutic strategies to modulate intestinal repair and adaptation.

## Footnotes

### Contributors

**Conceptualization**: AB, NB. **Methodology**: AB, NB. **Formal analysis**: AB. **Investigation**: AB. **Writing—original draft, review & editing**: AB, NB. **Visualization**: AB Supervision: NB. Project administration: NB. Funding acquisition: NB

### Funding

This research was supported by grants from NIH (R01AI148529 and R01AI148541 to NB) and NSF (IOS 2024252 to NB).

### Competing Interests

The Authors have no competing interests.

### Author Note

For AB, benchwork was done entirely at Cornell University, together with some data quantification. Additional quantifications, analysis, figure making and manuscript writing were done while at Zhejiang University.

## Acknowledgements

We thank Peter Nagy, George Samantsidis and Mikael Bjorklund for comments.

## AI use

ChatGPT was used midway through writing to polish language.

## Supplemental Material

This content has been supplied by the author(s). It has not been vetted by BMJ Publishing Group Limited (BMJ) and may not have been peer-reviewed. Any opinions or recommendations discussed are solely those of the author(s) and are not endorsed by BMJ. BMJ disclaims all liability and responsibility arising from any reliance placed on the content. Where the content includes any translated material, BMJ does not warrant the accuracy and reliability of the translations (including but not limited to local regulations, clinical guidelines, terminology, drug names and drug dosages), and is not responsible for any error and/or omissions arising from translation and adaptation or otherwise.

## Supplementary materials and methods

### Fly Stocks

*Drosophila melanogaster* stocks were maintained at room temperature (∼23°C) on yeast-cornmeal medium or at 18°C under a 12:12 hr light/dark cycle. Canton-S (Cs) (BDSC: 64349), was used as the wild-type control in all experiments not involving transgenic constructs. Gal4 drivers used included: ‘*w^-^; Esg-Gal4; UAS-GFP, tub-Gal80^TS^*’ (*Esg^TS^*, progenitor-specific) [10] and ‘*5966-GS*’ (EC-EB-specific, RU486-dependent gene-switch) [45]. UAS lines used were UAS-Histone2B-RFP [46], *UAS-rpr^OE^* (BDSC: 5823), and the *w^1118^* background strain.

### Experimental Design

All experiments were performed on mated female flies. Crosses and fly handling followed our previously described protocol [3], with minor adaptations. Briefly, conditional gene expression was induced using the TARGET system [47] or the GeneSwitch system (online supplemental figure 1A). For TARGET, flies developed at 18°C on a pre-experiment diet (yeast-cornmeal medium). F1 progenies were collected within 3 hours of eclosion and transferred to experimental diets. Transgenes were induced either by shifting to 29°C (TARGET) or by administering RU486 (100 μL of a 5 mg/mL solution in 80% ethanol added on top of food, with ethanol left to evaporate before adding flies) for GeneSwitch [48]. Genotype and treatment matched controls were used in all cases.

### Food Production

Standard, HS, and HY diets were prepared as previously described [3]. Briefly, agar was dissolved in boiling water, followed by addition of dry ingredients while constantly stirring. Acid mix was added at ∼60°C, and food was aliquoted into vials at ∼40°C.

### Bacterial cultures and oral infection

Bacterial cultures and oral infection were performed as previously described [18]. *Erwinia carotovora ssp. carotovora 15* (*Ecc15*)[49], its evf-deficient mutant (*Ecc15^evf-^*) [43] and *Escherichia coli* (*E.coli*) were maintained on standard LB agar. To obtain bacteria for infections, single colonies were inoculated into LB broth and grown at 29 °C, shaking for 16 hrs. Cultures were then pelleted (3,000 g, 5 min), and resuspended in PBS to an optical density at 600 nm (OD₆₀₀) of 200.

Adult flies were starved in empty vials for 2 h at 29 °C, then transferred to vials whose standard food surface was covered with a Whatman filter paper disk saturated with 150 µL of 2.5% sucrose and bacterial suspension (final OD₆₀₀ = 100). Flies remained at 29 °C until the designated dissection time point or survival assay endpoint.

### Immunochemistry

As previously performed [3], midguts were fixed in 4% paraformaldehyde, washed in PBS with 0.1% Triton X-100, and blocked in 1% BSA and 1% normal donkey serum. Immunostaining was performed using mouse anti-Dlg1 (1:100, DHSB 4F3, AB_528203) as primary antibody and Alexa Fluor-conjugated secondary antibodies (Thermo Fisher). DNA was stained with DAPI (1:50,000) and mounted in Citifluor AF1. Imaging was performed on a Zeiss LSM 700 confocal microscope.

### Posterior Midgut Area Measurements

Midgut area was quantified by capturing tiled 10× fluorescence images of whole midguts, assembled in Zen software (Zeiss). A custom FIJI [50] macro was developed and used to threshold midguts in a broad user-selected area containing the posterior region of the midgut. Binary masks were generated, holes filled, and area was quantified via particle analysis.

### Cell Size Measurement

EC area was manually measured in FIJI from anti-Dlg1, stained midguts imaged as Z-stacks with a 20× objective with Zen software (Zeiss). The largest visible plane of each cell was manually outlined in FIJI using the polygon tool. Approximately 30 adjacent cells per midgut were measured, as previously performed [3].

### Cell loss assay

For a visual scheme of this experiment, please refer to online supplemental figure 1A. *5966GS>Histone2B-RFP* flies developed on standard diet and were shifted at eclosion on either HS or HY diets for 5 days. Flies were then fed RU486 for 3 days, followed by 2 days without RU486. Flies on both diets were dissected before infection procedures (Time 0), starved for 2 hours, and kept on bacteria for eight hours. After infection, flies were shifted to either HS, HY or Agar (Ag, starvation) diets and dissected at 12, 24, 48, 96 and 240 hours post initial point of infection. UC flies were not subject to any treatment, and in case of change of diet, shifted from one diet to another directly.

Confocal Z-stacks (20×) of half midgut hemispheres were acquired and RFP+ and RFP− cell numbers were measured, together with the area of the midgut in the image, to calculate density of cells per midgut. This was multiplied for the total measure of midgut area. Cell density × area × 2 (to account for half midgut) gave total cells per region. Turnover was calculated as new cells divided by lost cells. Lost cells included both RFP+ and unmarked cells, accounting for incomplete labelling on day 0. All the calculations are as previously performed [3].

### Survival

For a scheme of this experiment, please refer to figure 6A. Canton-S flies developed on standard diet and were shifted at 29C from eclosion on HS diet for 5 days. Flies were then infected daily with either *Ecc15*, *Ecc15^evf^*^-^, or *E.Coli*. Flies were also kept in unchallenged conditions or infected once with *Ecc15*. Flies were infected for 8 hours and then shifted back to HS or to Ag diet for 16 hours before the next infection. Survival was checked daily. Only female flies were scored, but flies were cohoused with male flies to ensure mated status.

### Statistical analysis

Statistical analysis was performed as previously reported [3] using generalized linear mixed models using fitme from spaMM [51]. Briefly, repeats were included in the models as random effects. Models were tested for normal distribution with Shapiro– Wilk normality tests and homoscedasticity of the residuals with Brush–Pagan tests. Non normal or heteroskedastic samples were transformed to improve fit using either log, Box Cox or squared transformation (depending on what yielded samples both normal and homoscedastic). In case no transformations were able to eliminate heteroscedasticity (only for comparisons in figure 6), resid.model was added to the model to account for heteroscedasticity. Akaike information criterion was used to select the most performing model. Models were compared with ANOVA to infer significancy. Interaction between samples was also calculated with this method. To characterize differences multiple conditions, general linear hypotheses tests were applied, using a Tukey post hoc pairwise comparisons. Survival statistics were calculated via Cox proportional hazards mixed effects model using the coxme package in R. After performing ANOVA on the model including treatment vs the model not including treatment, Tukey post hoc test was run to identify significantly different treatment groups.

## Supplemental figure legends

**Online supplemental figure 1.**
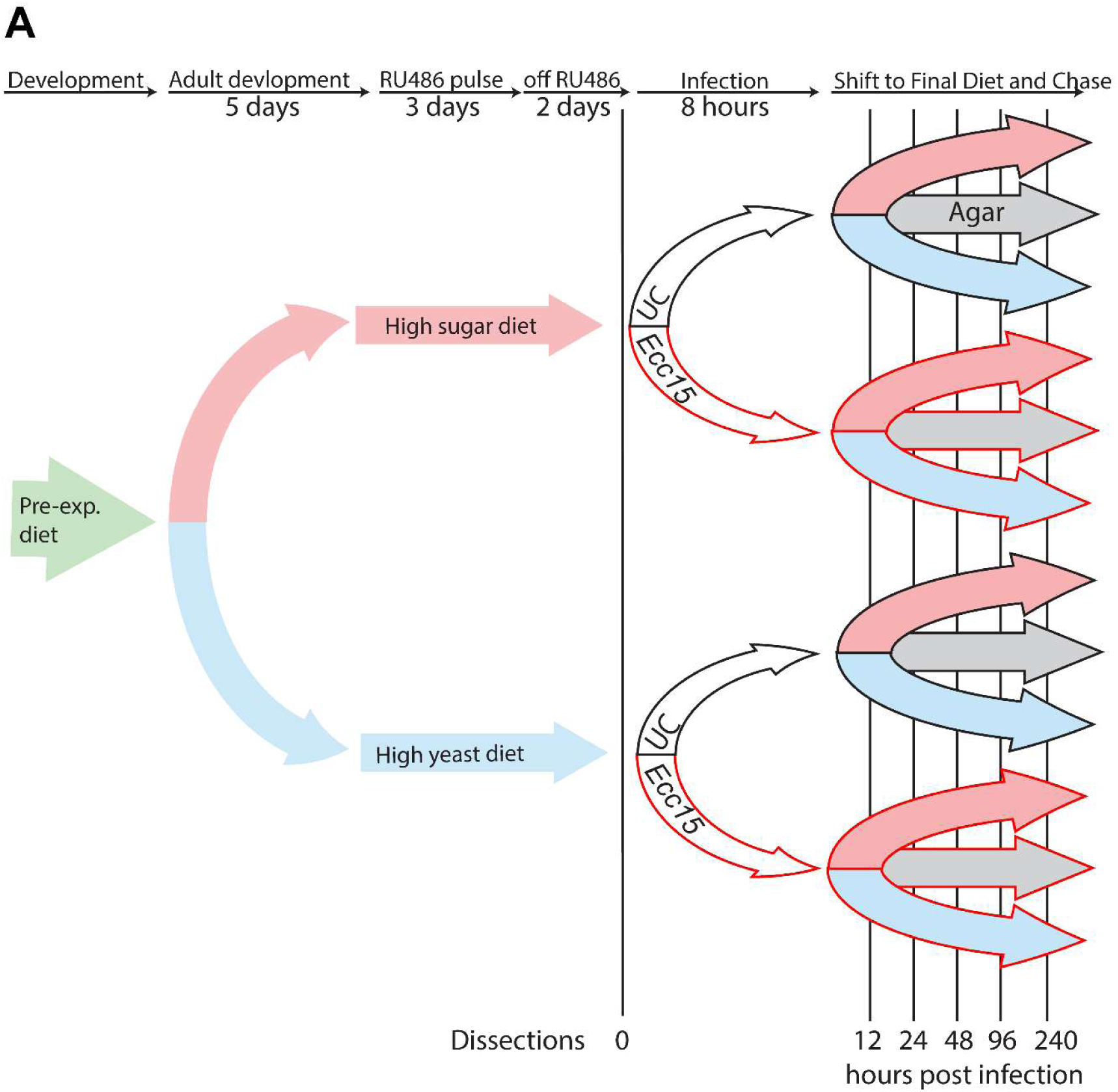
(A) Scheme depicting experimental strategy. Flies are kept during development on a pre-experimental diet at 25 °C. Upon eclosion, flies are transferred to either HS or HY diet at 29 °C for 5 days to allow for development. Flies are then transferred to the same diet but containing RU486 for 3 days, in order to activate the genetic system and mark ECs present at this point in time. Flies are then transferred to the same diet without RU486 for 2 days, to allow the drug to leave the flies. Flies are then either kept infected with *Ecc15* for 8 hours or kept unchallenged (UC) and then shifted to 1) the same diet (e.g. HS to HS), 2) the other diet (e.g. HS to HY) or 3) starved (e.g. HS to Ag). Flies were either dissected pre infection, or at 12,24,48,96 and 240 hours post infection.

**Online supplemental figure 2.**
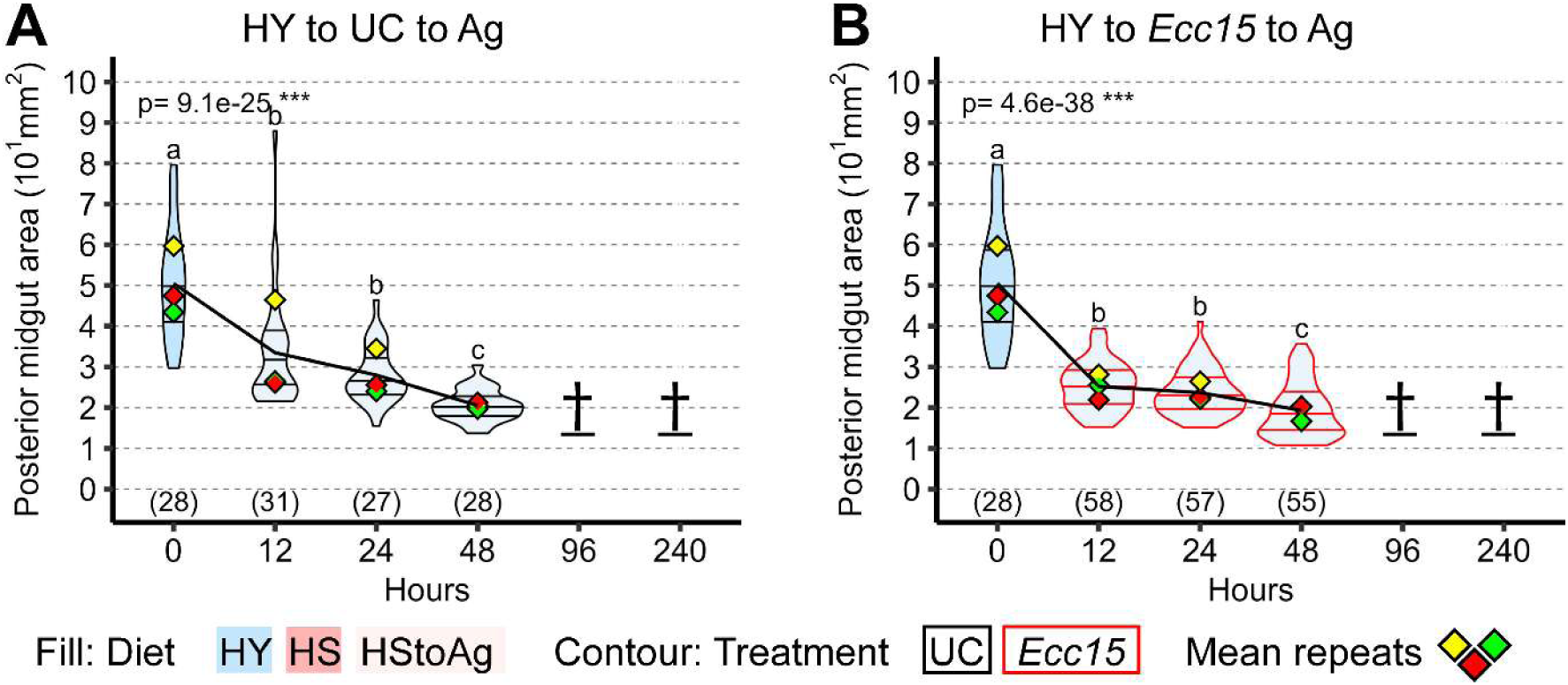
(A-B) Midguts shifted from high yeast (HY) diet to agar starvation diet (Ag) in Unchallenged Conditions (UC) progressively decreased in size (A). Upon infection, midguts shrink and then maintain this small size (B). For the violin/dot plots shown in this figure, white dots represent single posterior midgut area measurements. For the violin/dot plots shown in this figure, colored lozenges represent means of replicate experiments. Black line connects total means of each sample to show timeline of changes. Violin plot fills are color-coded according to diets (HS = red, HY = light blue, HY to HS = pink, HS to HY = purple) and their contour indicates treatment (UC = black, *Ecc15* = red). Numbers in parentheses at the bottom of charts indicate sample sizes. p indicates the result of ANOVA for samples in a single chart, and groups are obtained with Tukey Post Hoc test (letters above violins).

**Online supplemental figure 3.**
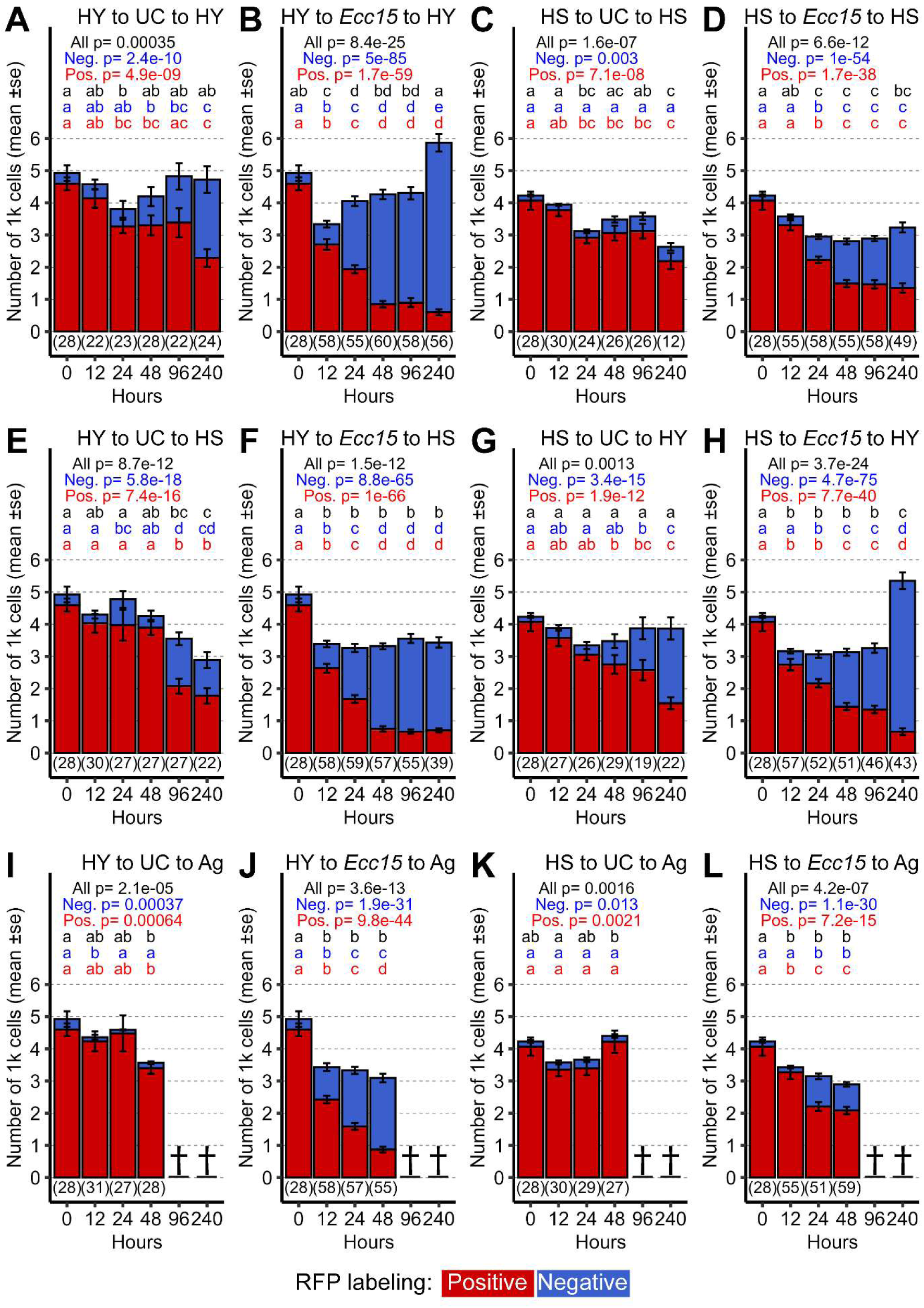
(A–L) Total cell number and turnover dynamics across all experimental conditions. (A) Flies maintained on a high-yeast (HY) diet throughout show only mild fluctuations in total cell number. These changes reflect moderate tissue turnover, with a progressive increase in the proportion of newly generated cells in the epithelium. (B) Infected flies kept on the HY diet before and after infection exhibit an initial decrease in total cell number, reaching a minimum at 12 hours post-infection. This is followed by marked regrowth, often surpassing pre-infection levels. (C) Flies maintained continuously on a high-sugar (HS) diet undergo gradual cell loss, with minimal production of new cells to compensate. (D) Infected flies kept on the HS diet before and after infection show more pronounced cell loss than uninfected controls, but this is partially offset by the production of new enterocytes, resulting in total cell numbers similar to unchallenged (UC) controls. (E) Flies initially maintained on HY diet and then shifted to HS diet display progressive cell loss. However, in contrast to flies constantly on HS (panel C), this loss is partially buffered by the production of new cells. (F) When HY-fed flies are shifted to HS diet following infection, total cell number drops sharply post-infection and then stabilizes. Despite this, tissue turnover remains elevated. (G) Flies initially on HS and then shifted to HY diet maintain stable total cell numbers and exhibit mild turnover, comparable to the HY-only condition in panel A. (H) Flies kept on HS prior to infection and shifted to HY post-infection display a mild drop in cell number, followed by substantial regrowth at 240 hours post-infection, along with elevated tissue turnover. (I) Flies initially on HY diet and then shifted to a starvation diet experience progressive cell loss with almost no turnover. All flies die by 48 hours post-treatment, the final recorded time point. (J) When HY-fed flies are shifted to starvation after infection, they exhibit a sharp initial drop in total cell number, accompanied by extensive tissue turnover. Again, no flies remain alive beyond 48 hours post-infection. (K) Flies initially kept on HS and then shifted to starvation maintain stable total cell numbers until succumbing to starvation. (L) Infected flies kept on HS and then moved to starvation show a slight reduction in total cell number and low levels of turnover before dying, with 48 hours post-infection being the last time point with surviving flies. All bar plots show RFP-positive (old) cells in red and RFP-negative (new) cells in blue, across time points. Statistical analyses compare treatments within the same condition for total cell number, RFP-positive cells (old), and RFP-negative cells (new).

**Online supplemental figure 4.**
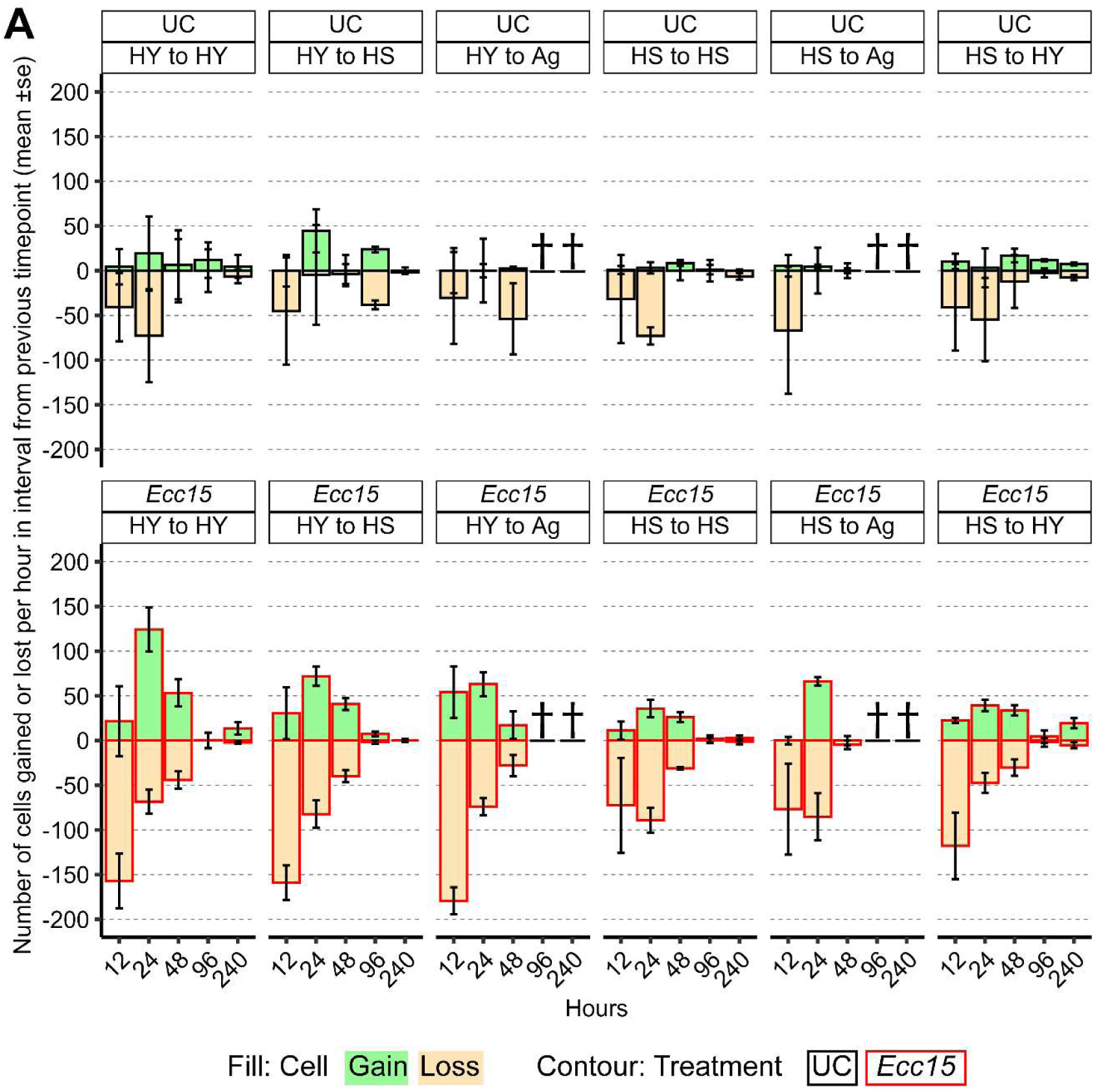
A) Number of cells gained or lost per hour, in intervals from previous timepoint, show that while unchallenged samples have a relatively flat distribution (top part of the chart), infection with *Ecc15* greatly induced cell loss at 12 hours for most conditions, then decreased gradually (bottom part of the chart). Cell gain follows closely the kinetic of loss, with a slight delay. All facets of the bar plot show cells lost per hour in yellow and gained in green, across time points. Bar plot contour indicates treatment (UC = black, *Ecc15* = red).

**Online supplemental figure 5.**
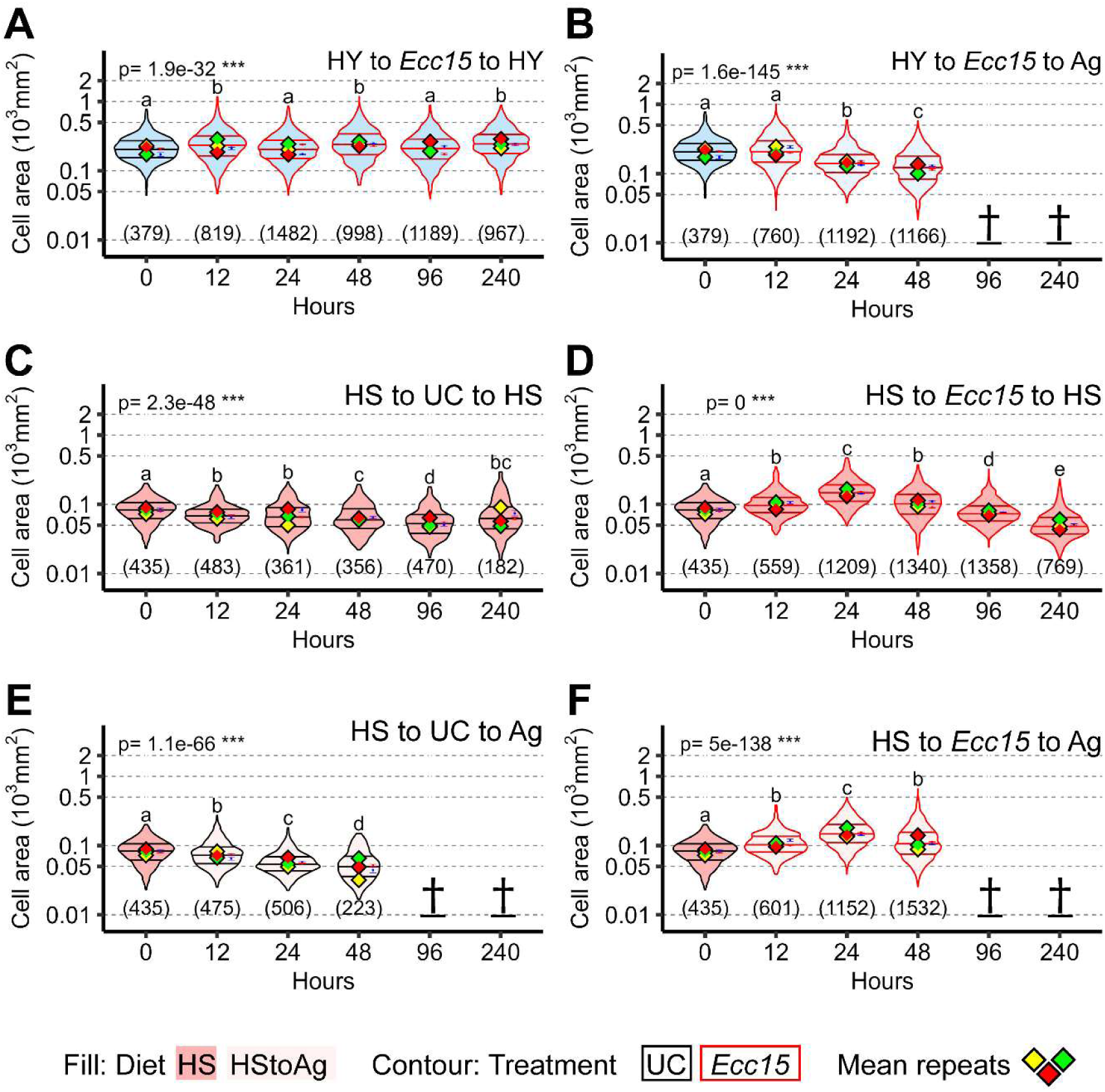
A-B) Infection does not reduce cell size in midguts starting from HY in several separate experiments. C-F) In midguts starting from HS diet (small) and kept on either HS or Ag after treatment, infection also does not lead to a shrinkage in cell size, but leads to a transitory increase (D, F), before eventually shrinking down due to the effect of diet, as visible also in UC samples (C, E). For the violin/dot plots shown in this figure, colored lozenges represent means of replicate experiments. Black line connects the total mean of each sample to show timeline of changes. Violin plot fills are color-coded according to diets (HS = red, HY = light blue, HY to HS = pink, HS to HY = purple) and their contour indicates treatment (UC = black, *Ecc15* = red). Numbers in parentheses at the bottom of charts indicate sample sizes. p indicates the result of ANOVA for samples in a single chart, and groups are obtained with Tukey Post Hoc test (letters above violins).

**Online supplemental figure 6.**
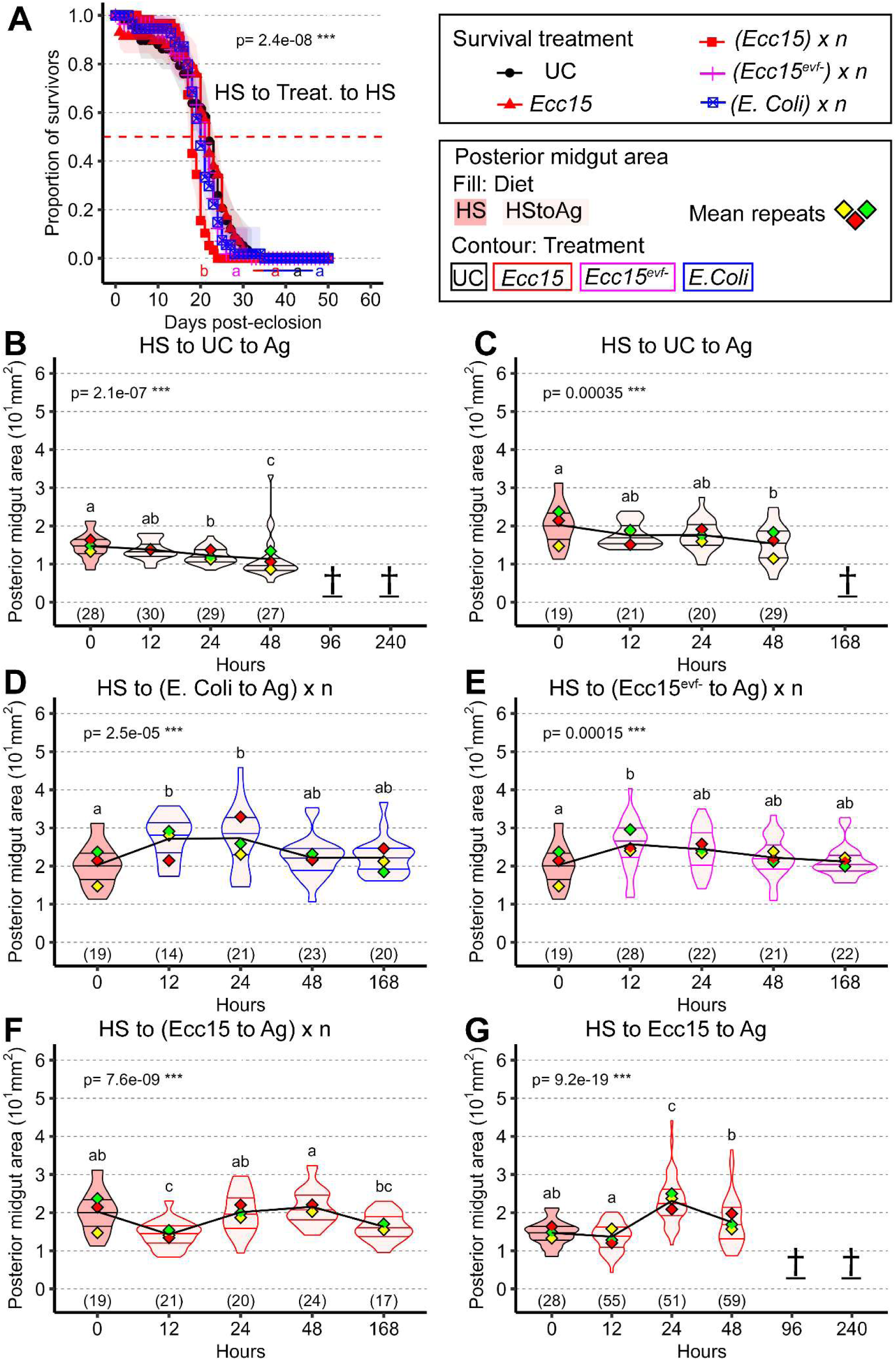
A) Survival for flies always kept on HS with different cyclical treatments shows that repeated infection with *Ecc15* in this case leads to a small decrease in survival compared to the rest of the samples. B-G) Midguts of flies shifted from HS diet to Ag keep a relatively small size, with further shrinkage upon prolonged stay on Ag diet and eventually early demise at 48 hours (B original panel of experiment, C control for microbial addiction). Similarly to samples always on HS, repeated addition of nonpathogenic microbes (*E.Coli*, D and *Ecc15^evf-^*, E) have an immediate effect in increasing the size of the midgut. Repeated infection with *Ecc15* in this condition does not lead to an increase in size of the midgut (F). If infected once with *Ecc15* before being shifted to Ag (G), flies show a transitory increase in midgut size at 24 hours post infection, followed by shrinkage and death in a similar manner as UC samples. This pattern repeated for a CantonS strain. For the violin/dot plots shown in this figure, colored lozenges represent means of replicate experiments. Black line connects total means of each sample to show timeline of changes. Violin plot fills are color-coded according to diets (HS = red, HY = light blue, HY to HS = pink, HS to HY = purple) and their contour indicates treatment (UC = black, *Ecc15* = red). Numbers in parentheses at the bottom of charts indicate sample sizes. p indicates the result of ANOVA for samples in a single chart, and groups are obtained with Tukey Post Hoc test (letters above violins). Survival statistics were calculated via Cox proportional hazards mixed effects model using the coxme package in R.

## Notes

### Competing Interest Statement

The authors have declared no competing interest.

